# Receive-Only Coupled Wireless Radiofrequency Probe for Endocavity MR Imaging Using a pTx System

**DOI:** 10.64898/2026.06.10.731354

**Authors:** Akbar Alipour, Volkan Acikel, Sayim Gokyar, Oktay Algin, Cagdas Oto, Priti Balchandani, Hilmi Volkan Demir, Ergin Atalar

**Affiliations:** BioMedical Engineering and Imaging Institute (BMEII), Icahn School of Medicine at Mount Sinai, New York, NY 10029; Aselsan, REHIS, 3051. Sok. No:3, 06830 Ankara, Turkey; GE HealthCare In, 9900 W Innovation Dr, Wauwatosa, WI 53226; Department of Radiology, Ankara University, Faculty of Medicine, Ankara 06800, Turkey; Faculty of Veterinary Medicine, Ankara University, Ankara 06110, Turkey; Department of Electrical and Electronics Engineering-National Magnetic Resonance Research Center (UMRAM)-National Nanotechnology Research Center and Institute of Material Science and Nanotechnology (UNAM)-Department of Physics, Bilkent University, Bilkent, Ankara 06800, Turkey; LUMINOUS! Center of Excellence for Semiconductor Lighting and Displays, School of Electrical and Electronic Engineering, School of Mathematical and Physical Sciences, School of Materials Science and Engineering, Nanyang Technological University, Singapore 639798, Singapore

**Author notes:** **Corresponding Author:** Akbar Alipour, MSc, PhD, JFISMRM. Assistant Professor, BioMedical Engineering and Imaging Institute (BMEII), Department of Diagnostic, Molecular And Interventional Radiology, Icahn School of Medicine at Mount Sinai, 3 East 101st Street, 7th Floor, Room 708, New York, 10029.

**Keywords:** Inductive coupling, interventional coil, passive RF resonator, parallel transmit system

## Abstract

**Purpose:** To enhance the SNR in MRI within a localized region of interest using a novel interventional wireless RF resonator probe combined with a dual-drive pTx system.

**Methods:** A dual-drive body birdcage coil was operated in a linearly-polarized mode to decouple a passive RF resonator probe from the transmit field while maintaining the resonator in a receive-only coupled mode. The resonator was fabricated using standard microfabrication techniques and tuned to the Larmor frequency of a 3T MRI system. The 10-g specific absorption rate (SAR) distribution was simulated to identify potential hot spots around the resonator prior to heating experiments. To evaluate the interaction between the resonator probe and the linearly-polarized transmit field, SNR and flip-angle distributions were measured in a phantom. In vivo imaging studies were subsequently performed using the resonator probe in con*j*unction with the linearly-polarized dual-drive birdcage coil.

**Results:** Temperature measurements demonstrated a normalized temperature increase of less than 0.10°C, corresponding to a SAR value below 1.21 W/kg. Experimental flip-angle mapping confirmed effective magnetic decoupling of the resonator probe using linearly-polarized dual-drive transmission. An SNR enhancement factor of 1.6 was achieved within the region of interest in phantom experiments. In vivo imaging demonstrated a 2.0 ± 0.2-fold SNR enhancement in the vicinity of the resonator probe.

**Conclusion:** A novel interventional approach for localized SNR enhancement in MRI was demonstrated using a wireless RF resonator probe and a dual-drive pTx system. The proposed technique enables local signal enhancement while minimizing transmit-field interactions, thereby facilitating safe interventional MRI and potentially improving image quality and diagnostic performance.

## INTRODUCTION

Endoscopy using alternative imaging modalities, including narrow-band imaging, magnifying endoscopy, optical coherence tomography, autofluorescence endoscopy, and intravascular ultrasonography, has been developed for the observation of suspicious lesions in the gastrointestinal tract and other narrow body orifices and endocavity regions [1-3]. However, poor tissue contrast and low SNR limit the utility of these techniques [4,5].

MRI, a non-ionizing imaging modality with high soft-tissue contrast, is widely used for imaging body orifices as well as for guiding minimally invasive interventions [6,7]. The MRI surface arrays used for imaging endocavity regions are the spine array, positioned beneath the patient, and the surface array, placed on the abdominal surface. These arrays are most sensitive at distances of 80-100 mm from the posterior and anterior subcutaneous fat layers, respectively. Given that the anterior-posterior dimension in adults typically ranges from 300-350 mm, a central region approximately 100-150 mm in diameter within body cavities (e.g., pelvis, prostate, and colon) remains imaged at a lower SNR. To address this limitation, various active and passive interventional MRI coils have been proposed to improve the SNR within a localized region of interest (ROI). Active coils usually rely on either an antenna or small coils connected to a separate receive channel [8-14]. This approach may pose safety concerns due to long cable connections, crosstalk, and potential malfunction of electronic components such as diodes and capacitors. In addition, cables are typically bulky and difficult to handle, and they can hinder the mechanical functionality of the imaging system. Nevertheless, this technique continues to be used in several MRI applications.

Another approach is the use of wireless passive radiofrequency (RF) coils, which are inductively coupled to the transmit and receive coils without any direct cable connection to the MR scanner [15,19]. This method is primarily used for interventional imaging and for tracking interventional devices. However, these coils or resonators are not decoupled from RF excitation; therefore, the transmit field induces current on the coil, resulting in an additional magnetic field that may distort the flip-angle distribution and cause heating. A possible solution is to decouple the passive coil from the transmit field using back-to-back diodes. However, these diodes are bulky and significantly increase the size of interventional coils, whose dimensions are often highly constrained.

Geometric decoupling is another method for decoupling passive coils from the transmit field using pTx systems, such as dual-drive birdcage coils. This can be achieved by steering the transmit field in a desired angular direction through either changing the feed location or simply rotating the linearly polarized birdcage coil. The same effect can also be achieved by ad*j*usting the excitation amplitudes of the input ports of a dual-drive birdcage coil [20-22].

pTx systems, such as dual-drive birdcage coils, have been introduced for tracking the position of interventional devices, detecting catheter rotational orientation, and reducing implant heating [23-25]. For instance, Eryaman et al. demonstrated a method for reducing RF heating of metallic devices using a dual-drive birdcage coil [23]. They showed that by controlling the excitation currents at the two ports of the birdcage coil, the RF current induced near the lead tip could be reduced to zero. In another study, Godinez et al. showed that safe operation of MRI-guided cardiovascular interventions using standard metallic guidewires is possible with pTx RF coils operated in a null-current mode, which minimizes induced RF currents and therefore enables safe tissue visualization [25].

In this study, by leveraging the dual-drive excitation technique, we introduce a method for interventional MR imaging that employs a new class of wireless RF resonator probes to improve SNR in the anatomy of interest. A passive RF resonator probe was fabricated using standard microfabrication and thin-film microwave techniques without the use of lumped-element components. A built-in distributed capacitor was used to tune the resonator for operation at 3T MRI systems. We employed a dual-drive transmit system to decouple the resonator from the transmit field while maintaining it in a receive-only coupled (Tx-decoupled) mode. The receive-only coupled resonator provided high-SNR signals in its vicinity without causing flip-angle distortion or RF heating.

This paper presents electromagnetic (EM) simulations performed to assess the electrical characteristics of the resonator, excitation field distributions, and specific absorption rate (SAR). It then describes the design and fabrication of the resonator probe prototype and concludes with validation of the proposed method in phantom and rabbit models.

## THEORY

Conventionally, a quadrature hybrid method is used in the standard birdcage coils to divide the excitation RF signal into two signals with 90° phase shift and equal magnitude to generate a circularly forward-polarized transmit field [20]. Considering a passive RF resonator inside a conventional forward-polarized birdcage coil (Fig. 1a), inductive coupling between the coil and the RF resonator leads to local MR signal enhancement in the vicinity of the resonator. The principle of local signal enhancement originated from two basic concepts: (i) transmit coupling: increased flip angle near the resonator during RF excitation; and (ii) receive coupling: the MRI signal enhancement during the reception due to the coupling between the transverse magnetization (*M*) and resonator [26-30].

**Figure 1:**
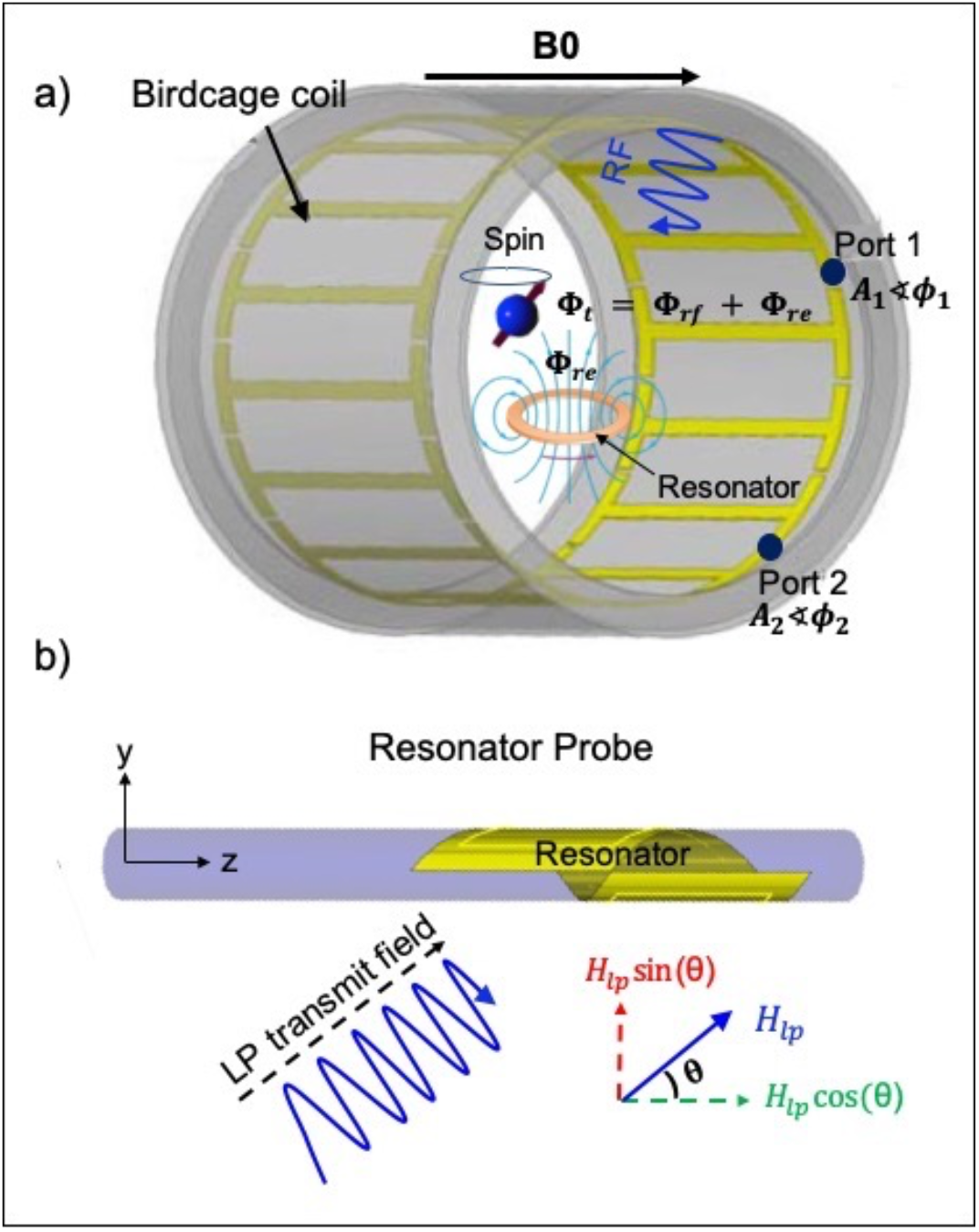
(a) Interaction of the circular forward-polarized magnetic flux (Ф_*rf*_) with a passive RF resonator. Inductive coupling between the resonator and the transmit magnetic flux results in an additional magnetic flux (Ф_*re*_) near ad*j*acent spins, therefore, total magnetic flux experienced by the ad*j*acent spins is Ф_*rf*_ + Ф_*re*_. This figure also shows a birdcage coil with dual-drive capability. Scaling the amplitude and phase at each port can generate different transmit polarizations. (b) The linearly-polarized transmit field (*H*_*lp*_) makes an angle *θ* with the surface normal vector (y-axis) of the resonator. *H*_*lp*_ can be decomposed into parallel and perpendicular components to the surface normal vector. The parallel component is the effective component that excites the spins and decouples the resonator.

The magnetic flux generated by the transmit birdcage coil (Ф*rf*) is inductively coupled to the resonator. The reaction of the resonator with Ф*rf* leads to circulating current in the resonator, which results in an additional magnetic flux (Ф*re*). Therefore, spins in the resonator vicinity experience a larger magnetic field (or flip angel) in comparison with other spins away from the resonator. If we consider the resonator as a simple series RLC circuit, the total flux (Ф) around the resonator can be written as:

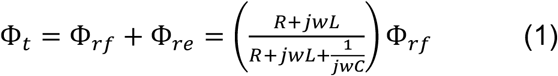

where, R, C, and L are the resistance, capacitance, and inductance values of the resonator, respectively.

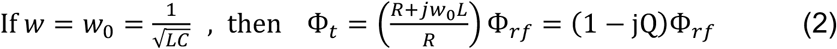

where *w* is the angular frequency, *w*_0_ is the angular Larmor frequency, *j* is the imaginary number defined by 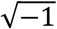 and represents a 90° phase shift between the transmit field and the field generated by resonator and Q = *w*_0_*L/R* is the resonator quality factor (Q-factor).

Flip angle amplification in the resonator vicinity depends on various parameters including the Q-factor, the orientation of the resonator, and the distance of the point where the flip angle is measured.

Considering a dual-drive linear polarized birdcage coil instead of a conventional quadrature hybrid one (Fig.1a). The EM fields generated by a dual-drive birdcage coil can be explained using the following equations [20]:

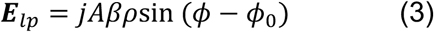

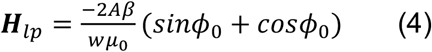

where ***E***_*lp*_ and ***H***_*lp*_ are the electric and magnetic fields generated by dual-drive birdcage coil, *A* is a constant depends on the excitation, *β* and *ρ* are parameters of Bessel function extracted from Maxwell equation solution in cylindrical coordinate, *ϕ* is the angular coordinate in the cylindrical coordinate system, and *ϕ*_0_ is the steering angle for the field pattern. The orientation of the linearly-polarized magnetic field generated by dual-drive birdcage coil can be steered in any angular direction by weighing the excitation currents of port 1 and port 2 with changing *ϕ*_0_.

If a dual-drive birdcage coil is fed with the same phase at port 1 and port 2, a linearly-polarized RF excitation can be obtained instead of a forward-polarized one. The coupling of the excitation field and the resonator is directly related to the applied excitation field orientation. Assuming an applied linearly-polarized transmit magnetic field (***H***_*lp*_) such that its excitation direction makes an angle *θ* with the surface normal vector of the resonator (Fig. 1b). Scaling the amplitudes at each port by changing *ϕ*_0_ allows steering of the linearly-polarized transmit field orientation (relative to the resonator probe), consequently changing the coupling level. ***H***_*lp*_ can be decomposed into two orthogonal components in z and y directions, respectively. ***H***_*lp*_ sin (θ) is the perpendicular component and ***H***_*lp*_ cos (θ) is the parallel component to the surface normal vector of the resonator. According to Faraday’s law of induction, only the parallel component can couple to the resonator. When the orientation of the linearly-polarized transmit field becomes perpendicular to the surface normal vector of the resonator probe, no coupling is expected. With this field, however, spin excitation is still possible. Therefore, the parallel component is the effective component that decouples the resonator from the transmit field, eliminates the current induction, and leaves the resonator in the receive-only coupling mode.

As it is clear from the E-field expression (equation 3), the field is suppressed on the whole *ϕ* − *ϕ*_0_ plane. If the resonator lies on the zero E-field plane, there will be no induced currents on the resonator. Setting the E-field to zero makes the parallel component of the H-field vanishes at the same plane; therefore, this technique prevents the inductive coupling between the transmit field and the resonator and leaves the resonator in receive-only coupled mode. This decoupling may prevent the possible high SAR hot spots and avoid the RF over-flipping in the vicinity of the resonator.

In the receive-only coupled mode, the rotating magnetization vector (*M*) generated by excited spins inductively couples to the resonator, resulting in an increased electromotive force (*ϵ*) induced in the receiver coil, which improves the receive signal sensitivity in the resonator vicinity.

If ***H***_*lp*_ is the magnetic field produced by the linearly-polarized birdcage coil, from the principle of reciprocity it can be shown that the incremental *ϵ* induced in the coil in the presence of the resonator [31]

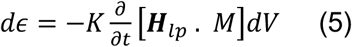

Integration over the volume, we drive the total MR signal (*t*)

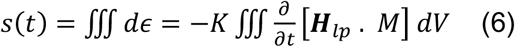

where *K* is a constant that depends on different parameters including the resonator Q-factor, the orientation of the resonator, and the distance of the point where the signal is measured.

Choosing the appropriate linearly-polarized dual-drive excitation angle, θ should set the resonator probe in receive-only coupled mode. By monitoring the flip angle variations in the resonator vicinity, the excitation pattern that satisfies the above conditions can be determined.

## METHODS

### RF Resonator Probe

The proposed passive RF resonator is a broadside-coupled multilayer structure with the upper and lower metal layers connected through metallized vias, with a dielectric layer sandwiched between them (Fig. 2a) [16,28]. The resonator was fabricated on a flexible dielectric substrate consisting of a 20-µm-thick polyimide film (Kapton® HN, DuPont, Berlin, Germany; ε_r_ = 3.7). A thermal deposition process was used to pattern 15-µm-thick gold layers on both sides of the Kapton substrate (Fig. 3a). A thickness of 15 µm was sufficient to prevent an increase in series resistance due to the skin effect, since the skin depth at 123 MHz is approximately 7 µm. A distributed capacitor formed between the two metal layers was utilized to create a series RLC resonant circuit tuned to the operating frequency of a 3T MRI scanner (123 MHz).

**Figure 2:**
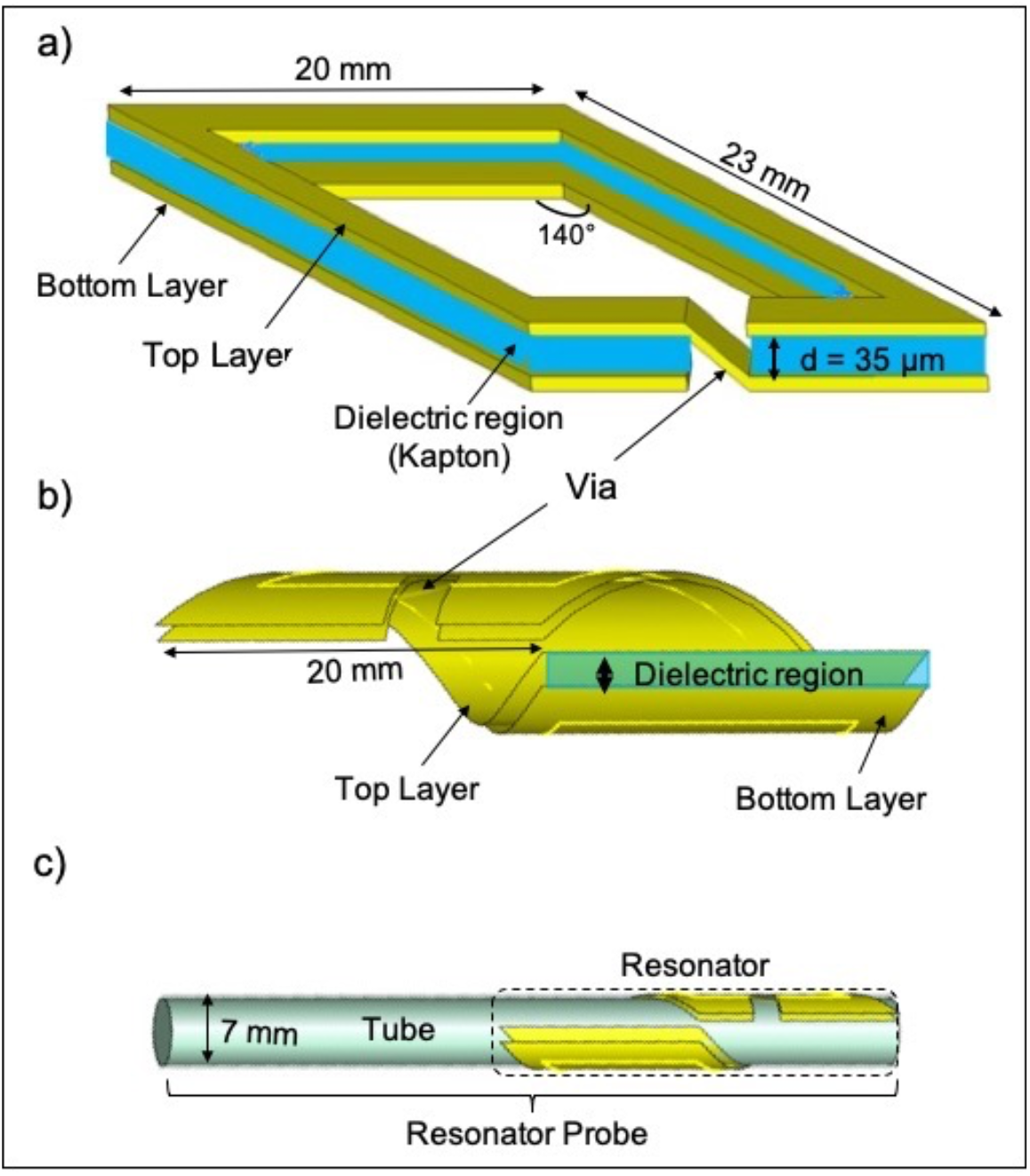
a) Schematic (not to scale) of the proposed flexible passive parallelogram RF resonator, which consists of two metal layers with a dielectric substrate (Kapton) sandwiched between metal layers. The metal layers are physically connected via metallization. b) The wrapped form of the resonator and specific geometry of the resonator preventing overlap. c) Schematic of the resonator probe. The resultant resonator was wrapped around polymer tube that was 7 mm in diameter.

**Figure 3:**
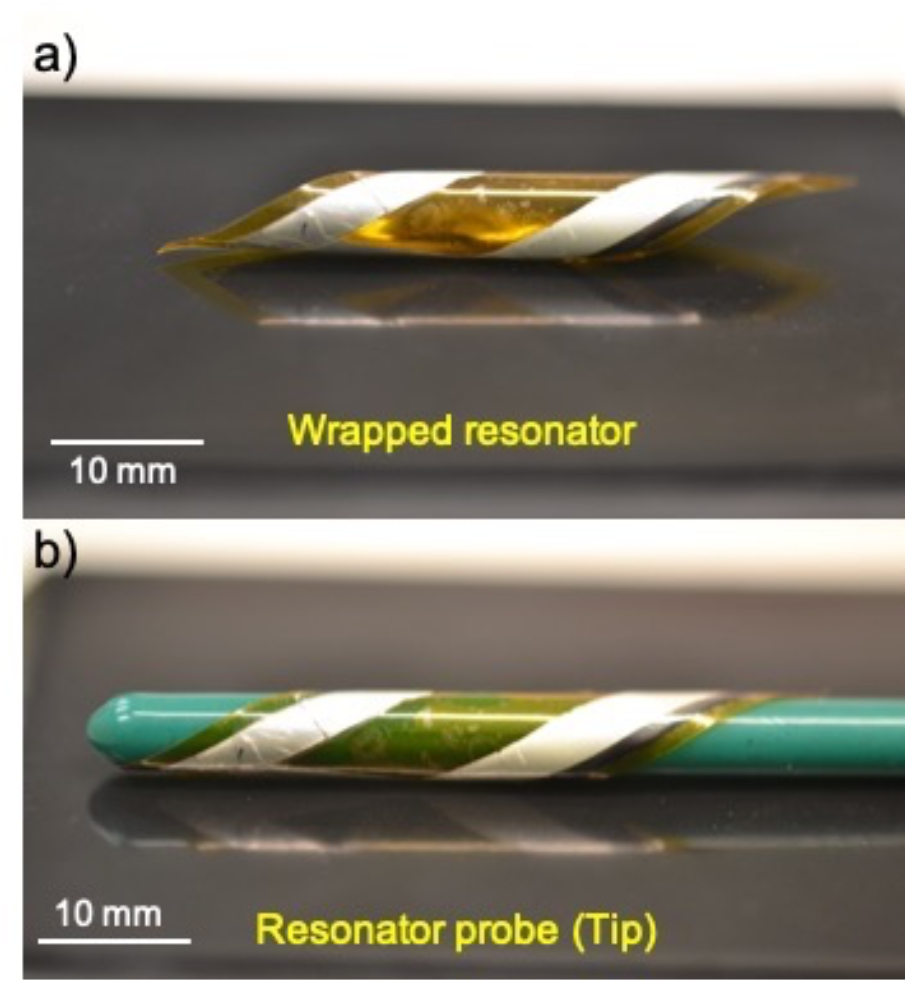
a) Photograph of the fabricated wrapped resonator. b) The resonator probe used in MR imaging. To ensure the device’s biocompatibility, the probe was coated with a thin layer of PDMS. This also facilitates the navigation of the probe inside the body.

The parallelogram geometry and flexible structure of the resonator allowed it to be easily wrapped around a cylindrical polymer tube with a diameter of 7 mm (Fig. 2b,c). Hereafter, the configuration shown in Fig. 2c is referred to as the “resonator probe.”

Finally, the resonator probe was coated with a 200-µm-thick layer of polydimethylsiloxane (PDMS) (Fig. 3b). PDMS (Sylgard® 184 Silicone Elastomer, Dow Corning, Sigma-Aldrich, Germany) is a widely used polymer in microelectromechanical systems and is extensively employed as a biocompatible material for implantable devices and structures [32]. The PDMS coating provides electrical insulation and biocompatibility while also reducing capacitive coupling between the resonator probe and the surrounding medium.

### Electromagnetic Simulations

Electromagnetic (EM) simulations were performed using CST Microwave Studio (Computer Simulation Technology, Darmstadt, Germany) to characterize the resonator design parameters and evaluate the transmit magnetic field 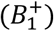 and local specific absorption rate (SAR) distribution in the presence of the resonator probe.

The dielectric thickness (*d*), which determines the capacitance value, is the primary design parameter controlling the electrical characteristics of the resonator, including its quality factor (Q-factor) and resonance frequency (*f*_0_). A series of EM simulations was conducted to investigate the effects of dielectric thickness on these electrical characteristics. The conductor length and geometry, which primarily determine the inductance of the resonator, were kept constant because the resonator was designed for a fixed probe diameter.

A body-sized birdcage coil operating in linearly polarized mode was modeled in CST Microwave Studio and loaded with a homogeneous phantom (conductivity, σ = 0.5 S/m; relative permittivity, ε_r_ = 80). Each port was tuned to 128 MHz and matched to 50 Ω. Simulations were performed with a convergence criterion of −30 dB using approximately six million mesh cells to calculate the 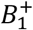 field and local SAR (W/kg) distributions in the presence of the resonator probe. The time-averaged 10-g SAR was calculated using the time derivative of the incremental electromagnetic energy. Regions exhibiting the highest SAR values in the vicinity of the probe were subsequently identified and used for experimental heating assessments.

### Bench Test

The resonator probe was immersed in a phantom (measured conductivity of σ=0.53 S/m and relative permittivity of ε_r_ = 75) and tuned with a pick-up coil connected to a network analyzer (Agilent E5061A, Santa Clara, California). The tuned *f*_0_ in the phantom tends to change when the probe is sited inside the body because the tissue loading is different from the loading in the phantom. Variable device placement limited us to an exact and quantitative estimation of the shifting values. Therefore, the *f*_0_ of the resonator probe was tuned roughly to the value experimentally predicted in the phantom.

### Phantom MRI Experiments

We performed a phantom experiment with an RF resonator probe to assess the proposed technique in a 3T MRI scanner (Trio, Siemens Healthineers) equipped with a dual-drive birdcage coil with using a phantom of 18 cm in diameter and 30 cm in length. The probe was placed in the phantom laying in the scanner z-direction. A dual-drive birdcage coil was used for transmission, and a 15-channel knee coil (Siemens Healthineers) was used for reception. First, feed ports of the dual-drive birdcage coil were calibrated by ad*j*usting the amplitudes and phases at each port.

Two feed points of the coil (outputs of the scanner amplifier) were connected to the oscilloscope through two equal-length coaxial cables. A gradient recall echo (GRE) sequence was run continually to allow a real-time RF signal calibration. The calibrated ports were connected to the dual-drive birdcage coil. As soon as the amplitudes and phases at each port were ad*j*usted, different polarization modes could be achieved using the dual-drive coil. For purpose of this study, the dual-drive birdcage coil was ad*j*usted to operate in the linearly-polarized mode by driving both ports in phase. We run the sequences manually using different linear excitation orientations with the dual-drive birdcage coil. The amplitudes of the applied voltage in port 1 and port 2 were weighted with cos*ϕ*_0_ and sin*ϕ*_0_ to control the EM field distribution inside the phantom. The value of *ϕ*_0_ was varied manually within the interval [0, *π*] with a step size of *π/*18. Scaling the amplitudes at each port allowed to steer the linearly-polarized transmit field orientation, consequently changed the coupling level. MRI signals were collected at various coupling levels using GRE sequences (TR = 12 ms, TE = 5.65 ms, FA = 10°, slice thickness 2 mm, matrix size = 256 × 256, field of view (FOV) = 134 × 134 mm^2^). For each image acquired at a specific coupling level, a circular ROI with a diameter of 3 cm was defined around the resonator probe for SNR measurements. SNR analysis was performed using the MRI signal images and separately acquired noise images obtained with the same pulse sequence parameters. The SNR was calculated according to the following equation [33]:

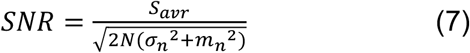

where *S*_*avr*_ is the averaged MR signal intensity; N is the number of array coils; and *σ*_*n*_ and *m*_*n*_ are the standard deviation and mean value of noise, respectively.

We also performed a resolution test using a phantom contained micro-bars of 150 µm in diameter and 300 µm periodicity. The resonator probe was positioned 3 mm above micro-bars. High-resolution MR images were obtained without and with Tx-decoupled (receive-only coupled) resonator using a GRE sequence with TR = 500 ms, TE = 8 ms, FA = 10°, slice thickness =1 mm, matrix size = 1024 × 1024, and FOV = 60 × 60 mm^2^.

To demonstrate the effectiveness of the method in transmit decoupling, flip angle maps were obtained in the phantom for both linearly-polarized and forward-polarized standard excitations using the double angle method [34].

### RF Safety Test

Heating tests were performed under linearly-polarized excitation in both Tx-decoupled and Tx-coupled modes to evaluate the effectiveness of the proposed method in reducing currents induced on the resonator probe. The experiments were conducted using a gel phantom during high-SAR imaging sequences at predetermined locations. The phantom was prepared according to the ASTM F2182 standard [35], with a conductivity of 0.47 S/m, a relative permittivity of 80, and a heat capacity of 4150 J/(kg·°C).

A high-SAR turbo spin-echo (TSE) pulse sequence (TR = 400 ms, TE = 15 ms, FA = 90°, FOV = 400 × 400 mm^2^, matrix size = 256 × 256) was applied continuously for 15 min. The resonator probe was positioned inside the phantom, and the phantom was centered within the magnet bore. The resonator probe was positioned at the location predicted by EM simulations to exhibit the highest electric field (E-field) intensity and, consequently, the greatest susceptibility to RF-induced heating [35].

Temperature measurements were obtained using fiber-optic temperature sensors connected to an optical thermometer (Neoptix RF-04-1 fiber-optic temperature sensor, Canada). The sensors, *p*_1_ − *p*_4_ were placed at potential SAR hot spots around the resonator probe, as identified through numerical simulations. A reference sensor *p*_5_ was positioned on the opposite side of the phantom. The temperatures measured at locations *p*_1_ − *p*_4_ were normalized to the temperature measured at the reference location, *p*_5_.

Following the ASTM standard test method F2182-11a, the actual SAR value around the resonator probe was calculated based on the heating measurements. SAR was calculated according to the equation:

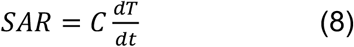

where C is the heat capacity of the gel phantom (4150 J/(kg.°C)). We calculated *dT*/*dt* using a linear fit over a 15-minute period. The SAR gain was calculated by dividing each calculated SAR by the reference SAR.

### In vivo Experiment

To evaluate the feasibility and effectiveness of the proposed MR imaging technique, an in vivo study was performed on a New Zealand White rabbit (2.8 kg) using a 3T MRI scanner. Prior to imaging, the animal was placed under general anesthesia and intubated. For premedication, atropine (1 mg/kg) was administered intramuscularly before anesthesia induction. General anesthesia was induced with intramuscular in*j*ections of xylazine (10 mg/kg) and ketamine (30 mg/kg). The animal experiments were approved by the Institutional Animal Care and Use Committee (IACUC) and were conducted in accordance with national guidelines and regulations for the care and use of laboratory animals.

The resonator probe was navigated through the rectum of the rabbit in the prone position. The linearly-polarized dual-drive birdcage body coil was used for transmission and a 15-channel knee coil was used for signal reception. A steady-state coherent sequences (FISP) with TR = 520 ms, TE = 4.18 ms, FA = 25°, matrix size = 384 × 384, FOV = 120 × 120 mm^2^, slice thickness = 1 mm were run for T_1_-weighted MR imaging and TR = 992 ms, TE = 46 ms, FA = 25°, matrix size = 256 × 256, FOV = 120 × 120 mm^2^, slice thickness = 1 mm for T_2_-weighted MR imaging. We used the same optimized linearly-polarized dual-drive transmission in the in vivo experiment as we used the phantom experiment. No additional coil optimization and calibration was performed for the in vivo study.

## RESULTS

### Electromagnetic Simulations

The numerically analyzed electrical characteristics of the resonator at different dielectric thicknesses (*d*) show that, as d increases, the distributed capacitance (*C*) decreases (Fig. 4a) because the separation between the two metal layers increases. Consequently, the resonance frequency shifts upward. In addition, larger values of d promote stronger localization of the electric field (E-field) within the dielectric layer, resulting in a higher Q-factor (Fig. 4b).

**Figure 4:**
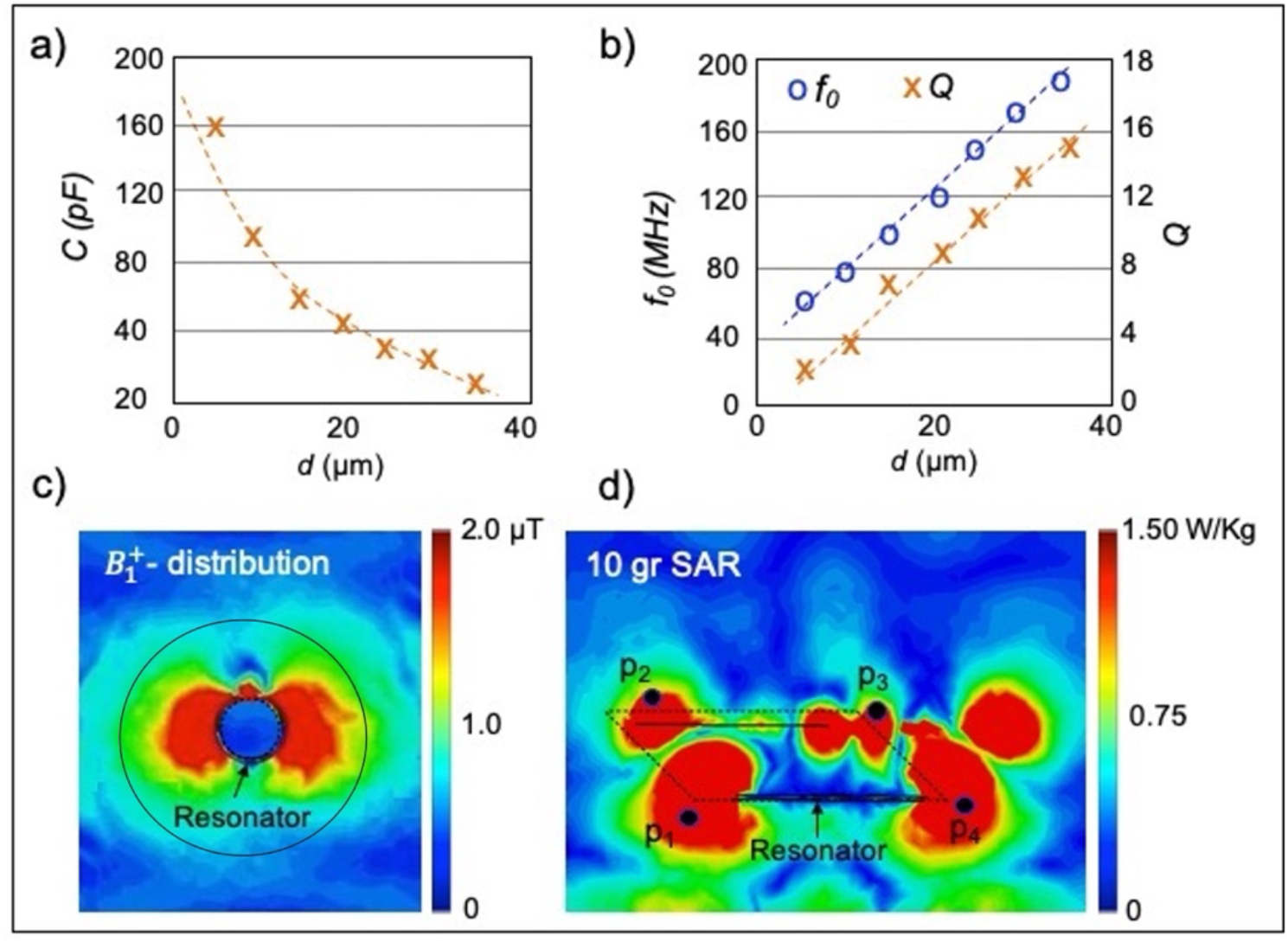
a) Simulation results detailing effects of the dielectric thickness (*d)* on the capacitance. As *d* increased, capacitance increased due to increasing the capacitive area. b) Simulation results detailing effects of *d* on the resonance frequency (*f*_0_) and Q-factor. *f*_0_ *and Q* increased as *d* increased. c) Simulated 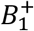 in the presence of the resonator probe shows that 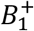 amplified in the vicinity of the resonator probe. d) transversal plane of a 10 gr SAR distribution around the coupled resonator. Edges of the resonator showed higher SAR values. Black dots (*p*_1_-*p*_4_) show the position of the temperature sensors, which were used for the heating test.

The total 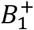 field generated by the transmit coil and the resonator probe is shown in Fig. 4c. The additional magnetic field produced by the resonator results in approximately a 3.3-fold enhancement of the 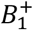 field within the ROI, indicated by the black circle.

The resulting maximum 10-g local SAR distribution of the resonator probe is shown in Fig. 4d. The peak local SAR was observed at the edges of the resonator and was approximately 2.4 times higher than that in other regions of the phantom.

### Phantom MRI Experiments

Figures 5a, 5b, and 5c show transverse phantom images acquired with the resonator probe under forward-polarized, reverse-polarized, and linearly polarized excitation, respectively. The inductive coupling between the forward-polarized transmit field and the resonator probe resulted in 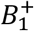 enhancement on one side of the resonator and cancellation on the opposite side (Fig. 5a). As expected, the reverse-polarized excitation did not directly excite the spins; however, it induced current in the resonator, generating a secondary linearly polarized magnetic field that excited spins in the vicinity of the probe (Fig. 5b). By setting *θ* to 0° (or 180°) through appropriate scaling of the excitation amplitudes at the input ports of the dual-drive birdcage coil, the generated linearly-polarized transmit field was aligned perpendicular to the surface normal vector of the resonator probe, resulting in transmit decoupling (Fig. 5c). The sensitivity enhancement observed around the circumference of the resonator probe was therefore attributable solely to receive coupling.

**Figure 5:**
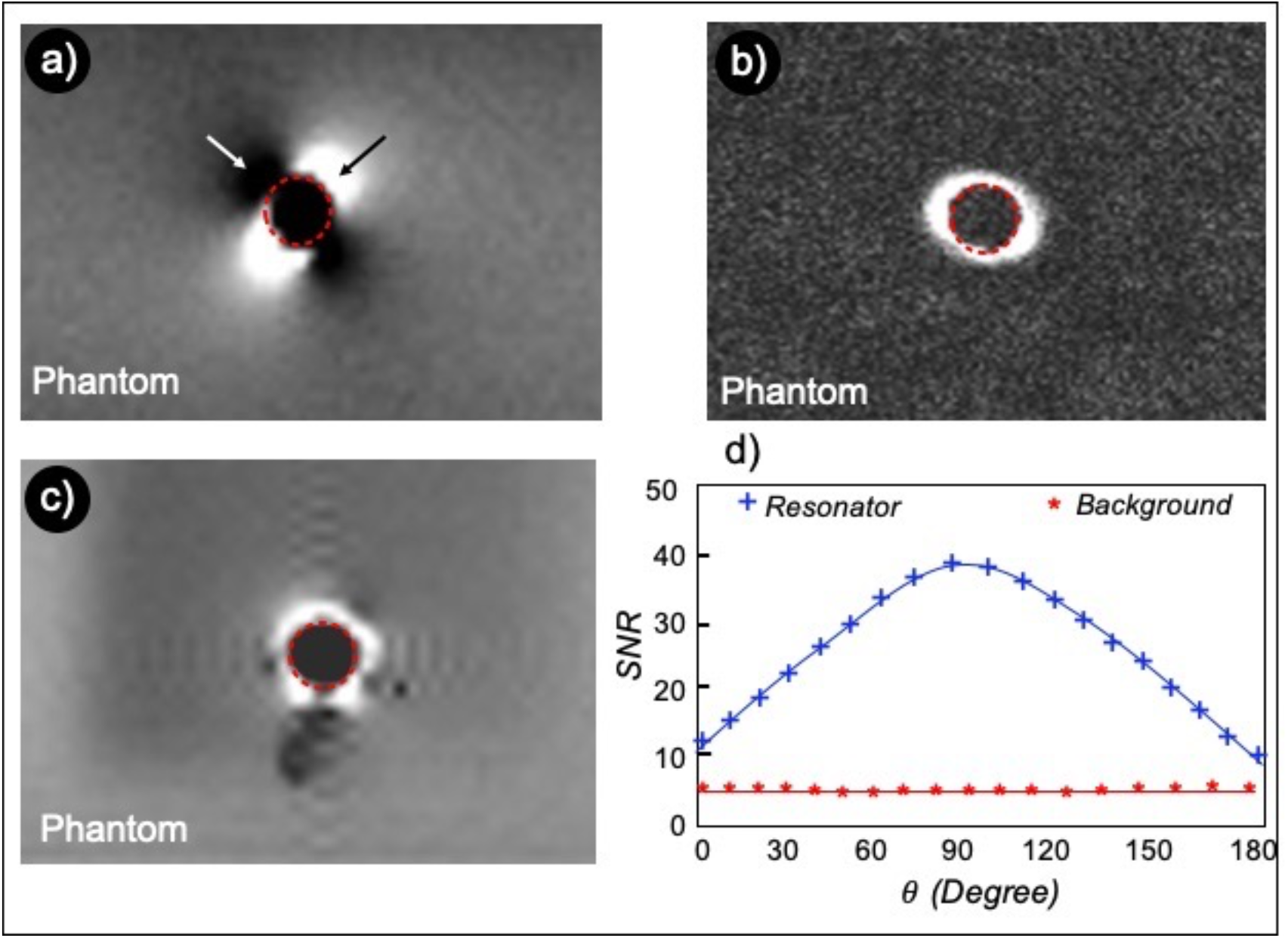
Transverse GRE images of the resonator probe (red dashed circle) inside the phantom were obtained using forward-polarized (a), reverse-polarized (b), and linearly-polarized (in receive-only coupled mode) (c) excitations. a) Forward-polarized transmit field excites the resonator and all spins inside the phantom. The additional magnetic field generated by the resonator resulted in RF amplification (black arrow) and cancellation (white arrow) in the resonator vicinity. b) Reverse-polarized excitation inductively coupled to the resonator probe and generated a secondary magnetic field, which only excited ad*j*acent spins. c) Linearly-polarized excitation is aligned to decouple the resonator probe. d) SNR values of the resonator probe and background phantom versus orientation of the linearly-polarized transmit field (*θ*). The background phantom for all excitations angles shows almost the same signal values. The coupling value between the transmit field and the resonator changes by *θ*. At *θ* = 0° and 180°, the resonator is in receive-only coupled (Tx-decoupled) mode.

Figure 5d shows the SNR measured within the resonator region and the background phantom at different coupling levels. The maximum SNR was obtained at *θ* = 90°, where the linearly-polarized transmit field was parallel to the surface normal vector of the resonator probe. As expected, the resonator was decoupled from the transmit field when the applied field was oriented perpendicular to the surface normal vector (*θ* = 0° and *θ* = 180°). The signal intensity of the background phantom remained unchanged as the transmit field orientation was varied.

High-resolution images (60 µm in-plane resolution) were acquired using a resolution phantom both in the presence and absence of the receive-only coupled resonator (Fig. 6a). The SNR enhancement in the immediate vicinity of the resonator probe improved the visibility of the micro-bars (Fig. 6b), which were not resolved in images acquired without the resonator (Fig. 6c). The presence of the resonator probe provided an average SNR enhancement factor of 1.6 within the ROI compared with imaging performed without the resonator.

**Figure 6:**
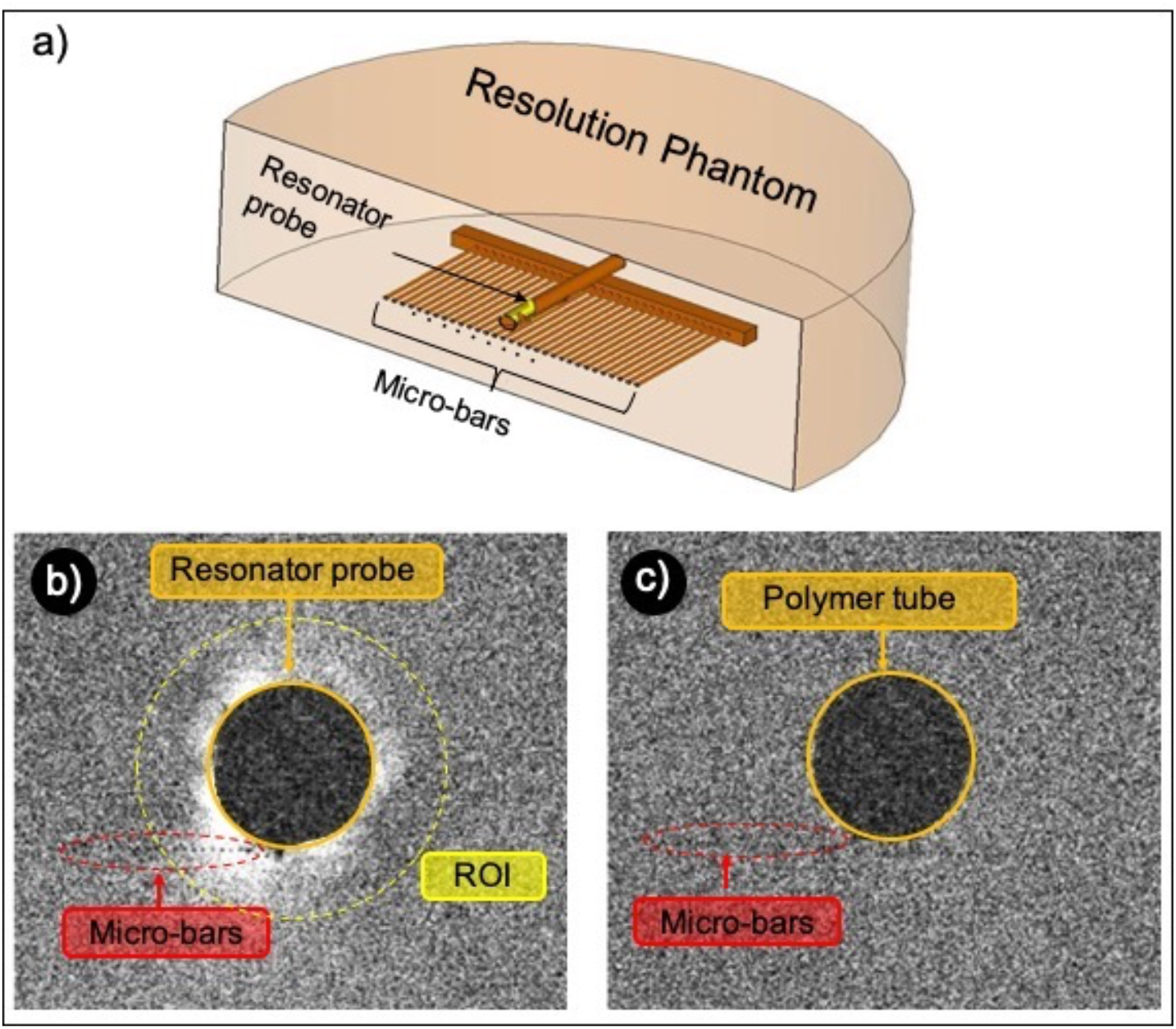
a) Schematic of the resolution phantom (not to scale). The phantom consists of fibers with 150 µm in diameter and periodicity of 300 µm. 60 µm in-plane high-resolution MR images obtained with the resonator probe in receive-only coupled mode (b) and without the resonator probe (c). SNR enhancement factor of 1.6 in the vicinity of the resonator allows for better micro-bars detection. Smaller pixel sizes resulted in lower SNR values in both images.

The pixel intensities of the 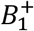 maps in the vicinity of the resonator probe as a function of the applied transmit voltage are shown for the maximum coupled mode (Fig. 7a) and the Tx-decoupled (receive-only coupled) mode (Fig. 7b) at various distances from the probe. The maximum coupling resulted in higher 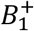 values near the resonator probe compared with the Tx-decoupled mode. At higher transmit voltages, the additional 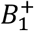 field generated by the resonator in the coupled mode caused RF over-flipping, resulting in a decrease in signal intensity in pixels ad*j*acent to the probe. In contrast, in the Tx-decoupled mode, increasing the transmit voltage led to a corresponding increase in signal intensity, as the resonator did not contribute to the transmit field and therefore did not induce local flip-angle amplification.

**Figure 7:**
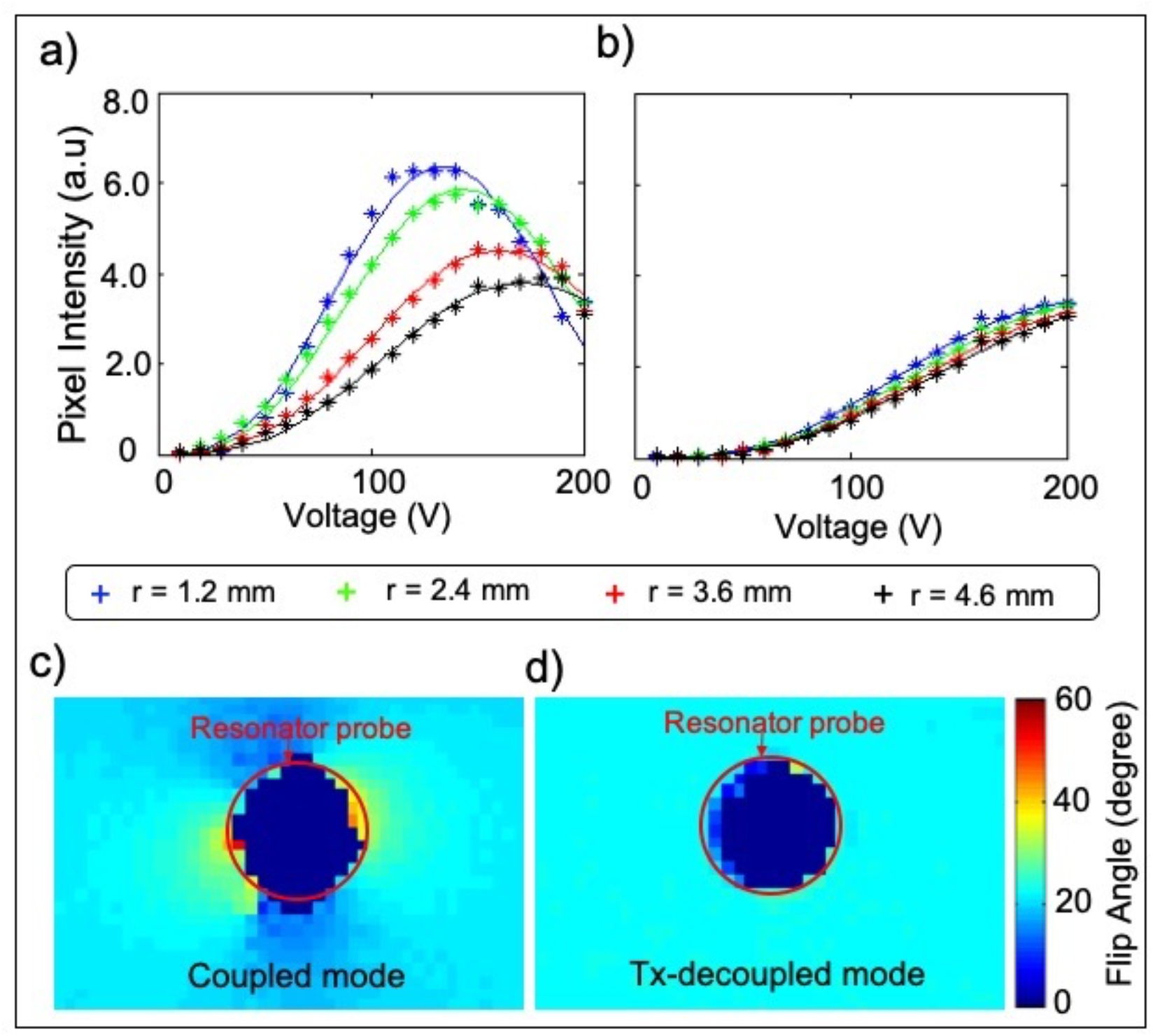
The pixel intensities of 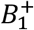 versus applied voltages were plotted at various distances from the resonator in the coupled (a) and Tx-decoupled (b) modes. a) A strong coupling between the resonator and linearly-polarized transmit field at higher voltages caused 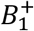 over-flipping. b) In the Tx-decoupled mode, 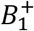 value was increase by increasing the voltage, independent of the distance. Flip angle maps were shown in the resonator vicinity in both coupled (a) and Tx-decoupled (b) modes. In coupled mode, induced current on the resonator resulted in flip angle amplification. A negligible flip angle distortion was displayed during the Tx-decoupled mode.

The flip-angle maps around the resonator probe in both the maximum coupled mode (*θ* = 90°) and the Tx-decoupled mode (*θ* = 0° and *θ* = 180°) are shown in Figs. 7c and 7d, respectively. In the coupled mode, the resonator probe amplified the local flip angle, resulting in flip-angle inhomogeneity in its vicinity. In contrast, only negligible distortion of the flip-angle distribution was observed in the Tx-decoupled mode.

### RF Safety Test

Prior to the heating experiments, EM simulations were performed to identify potential SAR hot spots around the resonator probe (Fig. 4d). Temperature measurements obtained at predetermined locations around the resonator probe showed a maximum normalized temperature increase of 1.41°C at point *p*_1_ in the coupled mode (Fig. 8a). The corresponding SAR value at this location was estimated to be 3.81 W/kg. This temperature increase was substantially reduced using the linearly polarized decoupling technique (Fig. 8b), which effectively decoupled the resonator probe from the transmit field. Using this approach, the maximum normalized temperature increase and corresponding SAR value at the same location *p*_1_ were reduced to 0.11°C and 1.21 W/kg, respectively.

**Figure 8:**
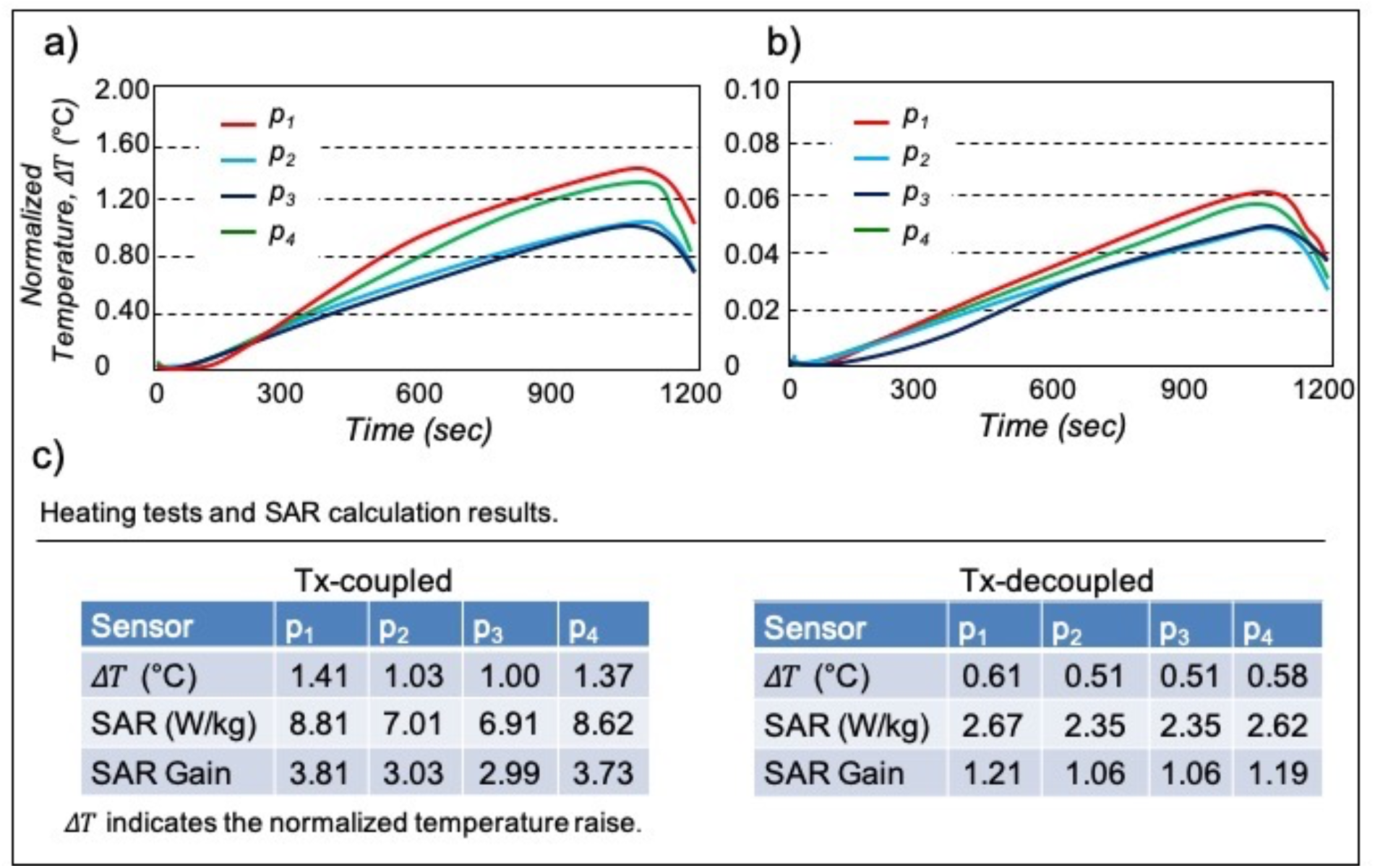
a) Heating tests were performed under high SAR sequences in both coupled (a) and Tx-decoupled (b) modes under linearly-polarized transmission. Four temperature sensors (*p*_1_-*p*_4_) were positioned at different spots around the resonator probe and the reference temperature was collected using sensor *p*_5_. In coupling mode, a maximum temperature increase of approximately 1.41°C was recorded at *p*_1_. This value decreased to 0.11°C in Tx-decoupled mode. c) Summarized temperature increase (Δ*T*), SAR, and SAR gain. Using the decoupling method, maximum SAR gain was mitigated by a factor of 3.3.

Figure 8c summarizes the measured temperature increases and corresponding SAR values at different locations along the resonator probe. The reference location, *p*_5_, exhibited a temperature increase of 0.5°C, corresponding to an SAR value of 2.31 W/kg.

### In vivo Imaging

T_1_-weighted sagittal and T_2_-weighted axial images were acquired in the Tx-decoupled (receive-only coupled) mode with the resonator probe positioned ad*j*acent to the colon wall, enabling SNR enhancement of colon wall structures in the vicinity of the probe (Figs. 9a and 9b). The acquired images demonstrated a 2.0 ± 0.2-fold SNR enhancement within colon-wall regions located 0.5-4.0 mm from the resonator probe. A reformatted axial image generated from the region indicated by the yellow dashed circle in Fig. 9a demonstrates the high level of anatomical detail that can be achieved in the endorectal region (Fig. 9c). A zoomed view of the resonator probe and its surrounding tissue from Figure. 9b further highlights the enhanced imaging region.

**Figure 9:**
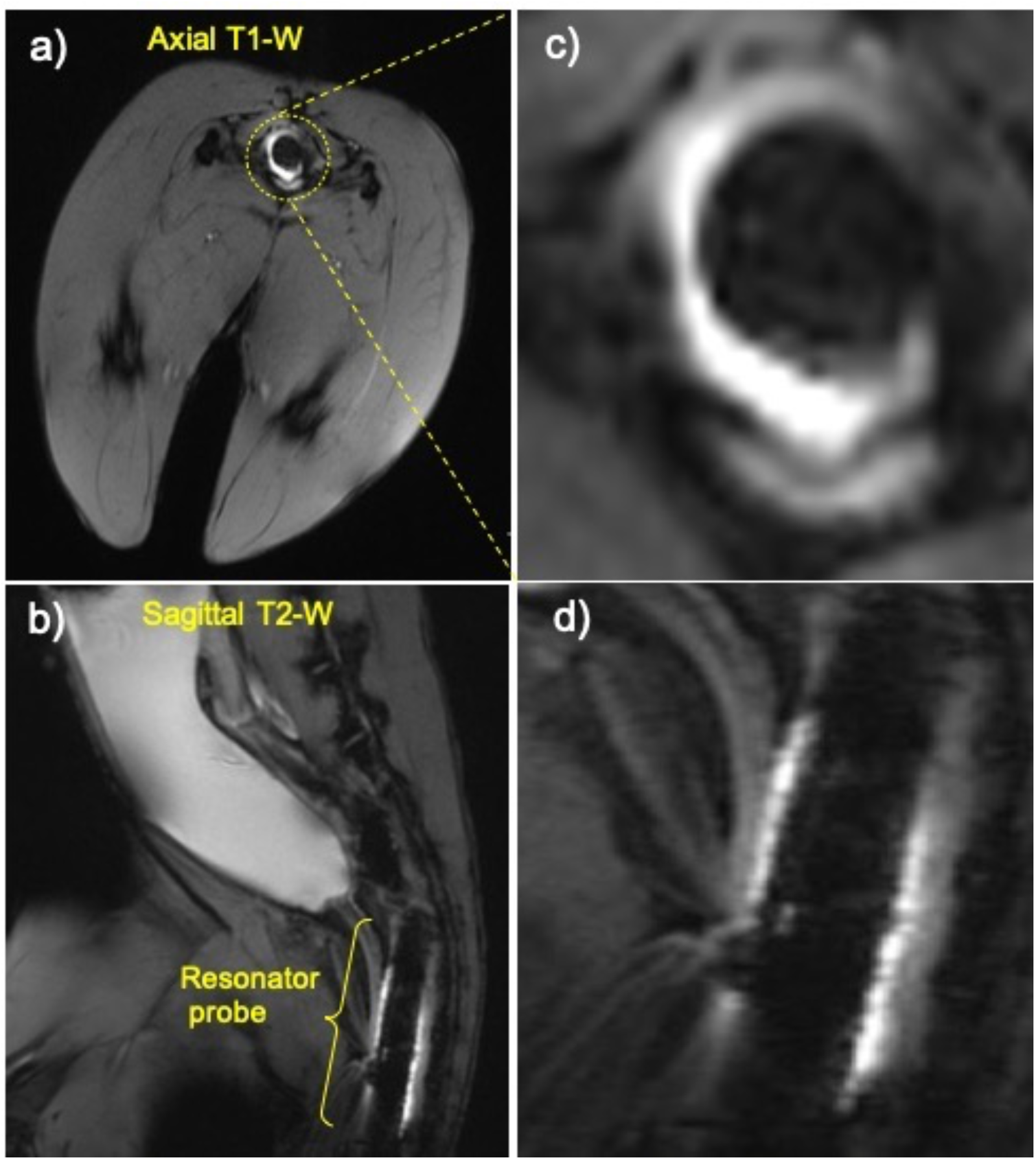
Proof of concept in vivo images obtained in a rabbit using the resonator probe in con*j*unction with the linearly-polarized dual-drive transmission. T_1_-weighted axial (a) T_2_-weighted sagittal (b) images demonstrate the image quality achievable in the region around the resonator probe. c) A 1 mm slice-width reformatted axial image taken from the dashed-yellow circular region of ***a***. d) Reformatted sagittal image taken from the region around the resonator probe of ***b***.

## DISSCUSION and CONCLUSION

A passive RF resonator probe was used in combination with a linearly polarized dual-drive coil to obtain SNR-enhanced images of a localized ROI in interventional MRI. Preliminary phantom and in vivo experiments demonstrated that the proposed RF excitation technique improved SNR in the localized region in the presence of the resonator probe without causing RF over-flipping or heating. An SNR enhancement of approximately 2.0 ± 0.2-fold was achieved at distances of 0.5-4.0 mm from the resonator probe surface in the in vivo experiments.

The resonator introduced in this work is a passive RF structure that does not require elongated wires for connection to the scanner [26-30]. It features a flexible construction and relies exclusively on distributed components to operate at the resonance frequency of a 3T MRI scanner. This design enables straightforward modification and implementation for use in various endocavitary regions, including the colon, vagina, prostate, and esophagus.

In this study, the conventionally used circularly forward-polarized excitation was replaced with linearly polarized excitation to cancel the transmit coupling between the resonator probe and the RF excitation field, thereby maintaining the resonator in a receive-only coupled mode via a pTx system. The two ports of the dual-drive birdcage coil were driven in-phase, and the amplitude at each port was scaled to steer the orientation of the linearly polarized transmit field [20–22]. The degree of Tx-decoupling between the linearly polarized transmit field and the resonator probe depends on the transmit field orientation relative to the resonator. When the applied transmit field was oriented perpendicular to the surface normal vector of the resonator probe, transmit coupling was canceled and only receive coupling remained. Consequently, signal amplification during reception was the sole contributor to the improved SNR in the vicinity of the resonator probe.

Experimental SAR calculations revealed a maximum SAR gain of 3.81 W/kg near the resonator in Tx-coupled mode, which the proposed decoupling method reduced to 1.21 W/kg. The Tx-decoupled resonator exhibited lower SAR gain owing to the intrinsic decoupling of the electric field from the resonator probe.

To demonstrate the efficacy of the probe in a relevant animal model, in vivo experiments were performed in rabbits, enabling data collection to assess safety and effectiveness prior to eventual translation to human use. Due to the restricted dimensions of the rabbit anatomy, the maximum diameter of the resonator probe was limited to 7 mm, which constrained the anatomical coverage within the ROI. The device can be readily scaled to extend anatomical coverage for human applications.

The *in vivo* experiment served as a proof-of-concept evaluation of the resonator probe performance in a rabbit model. Beyond this initial demonstration, the proposed method can be integrated into endoscopic probes to facilitate MR-guided navigation through complex anatomies, such as the gastrointestinal tract, enabling high-resolution imaging of the tract walls.

The use of linearly polarized dual-drive excitation preserves the prescribed flip angle distribution and mitigates heating, the principal concerns associated with standard forward-polarized excitation in the presence of a passive RF resonator. However, linearly-polarized excitation is less efficient from a global SAR perspective, as it comprises both forward and reverse polarized field components, effectively doubling the whole-volume average SAR per unit flip angle. Additionally, dual-drive excitation may reduce RF field homogeneity, particularly at high field strengths. The proposed technique could be adapted to operate with parallel transmission systems to leverage the increased degrees of freedom afforded by multiple transmission elements.

This interventional technique may enable improved visualization of fine anatomical features, thereby benefiting both image-guided interventions and diagnostic performance. Future work will focus on scaling the resonator design for human applications and evaluating its use in combination with linearly polarized transmission for imaging of the prostate and colon.

## ACKNOWLEDGMENTS

Authors gratefully acknowledge Berk Silemek, Umut Gundogdu, and Mustafa Utkur for their valuable discussions. The authors also thank Mustafa Delikanli for his technical support.

## Notes

### Competing Interest Statement

The authors have declared no competing interest.

